# Improving Landsat predictions of rangeland fractional cover with multitask learning and uncertainty

**DOI:** 10.1101/2020.06.10.142489

**Authors:** Brady W. Allred, Brandon T. Bestelmeyer, Chad S. Boyd, Christopher Brown, Kirk W. Davies, Lisa M. Ellsworth, Tyler A. Erickson, Samuel D. Fuhlendorf, Timothy V. Griffiths, Vincent Jansen, Matthew O. Jones, Jason Karl, Jeremy D. Maestas, Jonathan J. Maynard, Sarah E. McCord, David E. Naugle, Heath D. Starns, Dirac Twidwell, Daniel R. Uden

## Abstract

1. Operational satellite remote sensing products are transforming rangeland management and science. Advancements in computation, data storage, and processing have removed barriers that previously blocked or hindered the development and use of remote sensing products. When combined with local data and knowledge, remote sensing products can inform decision making at multiple scales.
2. We used temporal convolutional networks to produce a fractional cover product that spans western United States rangelands. We trained the model with 52,012 on-the-ground vegetation plots to simultaneously predict fractional cover for annual forbs and grasses, perennial forbs and grasses, shrubs, trees, litter, and bare ground. To assist interpretation and to provide a measure of prediction confidence, we also produced spatiotemporal-explicit, pixel-level estimates of uncertainty. We evaluated the model with 5,780 on-the-ground vegetation plots removed from the training data.
3. Model evaluation averaged 6.3% mean absolute error and 9.6% root mean squared error. Evaluation with additional datasets that were not part of the training dataset, and that varied in geographic range, method of collection, scope, and size, revealed similar metrics. Model performance increased across all functional groups compared to the previously produced fractional product.
4. The advancements achieved with the new rangeland fractional cover product expand the management toolbox with improved predictions of fractional cover and pixel-level uncertainty. The new product is available on the Rangeland Analysis Platform (https://rangelands.app/), an interactive web application that tracks rangeland vegetation through time. This product is intended to be used alongside local on-the-ground data, expert knowledge, land use history, scientific literature, and other sources of information when making interpretations. When being used to inform decision-making, remotely sensed products should be evaluated and utilized according to the context of the decision and not be used in isolation.

## 1 Introduction

The ability to monitor rangeland vegetation and to quantify changes in cover with satellite remote sensing is revolutionary to the rangeland management discipline. Whereas on-the-ground data collection and monitoring is constrained logistically, satellite remote sensing scales easily, measuring the landscape across space and through time. Satellite measurements are modeled to predict rangeland indicators, providing key information for land managers and practitioners globally (Hill & Guerschman, 2020). Chief among these indicators is vegetation cover at species or functional group levels. Historically, cover was broadly categorical or thematic, and occurred at local, regional, or national levels (Homer et al., 2015). More recently, fractional cover is used to preserve the inherent complexity and heterogeneity of the landscape, estimating the proportion of an area covered by vegetation or land cover type (Xian, Homer, Rigge, Shi, & Meyer, 2015). Fractional cover predictions, combined with local data and knowledge, can inform decision making at multiple scales, providing land managers flexibility that is absent with categorical classifications and severely lacking with on-the-ground observations (Kennedy et al., 2014).

Fractional cover products are widely available for United States rangelands (Jones et al., 2018; Zhang, Okin, & Zhou, 2019; Rigge et al., 2020). To produce such products, on-the-ground data is correlated to remotely sensed measurements using regression tree approaches, with models developed and predictions performed individually for each desired component. Although robust, univariate regression trees do not capitalize on the ability to learn from shared representation among dependent variables. That is, relationships among functional groups are not learned and therefore may not be reflected in predictions. A learned multitask model, however, examines all output variables together and learns from inherent interactions and relationships present in the data, improving learning efficiency and prediction accuracy (Caruana, 1997). Variables that covary will be reflected in the model, e.g., functional groups or species that are mutually exclusive or inversely related.

As these fractional cover products are the predictions of models, error and uncertainty are always present (Foody & Atkinson, 2003). Prediction error is most commonly calculated as the difference between a single on-the-ground measurement and its predicted value, a calculation that can be done with certainty. Multiple errors can then be averaged to produce a generalized accuracy of the model, e.g., root mean square error. Prediction uncertainty, however, differs from error in that it represents prediction confidence, or an expression of what is not known (Kendall & Gal, 2017). Consider the widely used example of a model that predicts whether an object in an image is a cat or dog, and is trained using only cat and dog data: if given cat or dog data, prediction confidence should be high; if given penguin data, prediction confidence should be low, as the model is unfamiliar with penguin data. While prediction error can only be calculated using individual on-the-ground measurements–and then averaged to obtain a generalized model accuracy–uncertainty can be calculated with every prediction, increasing its spatiotemporal utility. For the practitioner, uncertainty information can be used to assess prediction confidence, reliability, or use.

We describe a new rangeland fractional cover product that spans the western United States. We build upon previous advancements (Jones et al., 2018) by 1) utilizing a learned multitask approach to model the dynamic interactions of functional groups; and 2) generating pixel-level estimates of prediction uncertainty. We produce the fractional cover product annually from 1984 to 2019 at a moderate resolution of 30m. It is made available for analysis, download, and visualization through the Rangeland Analysis Platform (https://rangelands.app/) web application.

## 2 Materials and Methods

### 2.1 Data

#### 2.1.1. Rangeland Analysis Platform - fractional cover datasets

Jones et al. (2018) (hereafter referred to as fractional cover version 1.0) described the initial model and product released on the Rangeland Analysis Platform in 2018. The new model and subsequent product described in this paper (hereafter referred to as fractional cover version 2.0) supersedes the initial version.

#### 2.1.2 Vegetation field data

We used 57,792 vegetation field data plots collected by the Bureau of Land Management Assessment, Inventory, and Monitoring and Landscape Monitoring Framework, and the Natural Resources Conservation Service National Resources Inventory programs (Nusser & Goebel, 1997; Toevs et al., 2011). Plots were collected from 2004 to 2018 across uncultivated, undeveloped, privately and publicly owned rangelands across 17 western US states. We followed methods outlined in Jones et al. (2018), who aggregated species into the following functional groups: annual forbs and grasses, perennial forbs and grasses, shrubs, trees, litter, and bare ground. We randomly (stratified by state) divided the vegetation field data into training (90%, 52,012 field plots) and validation (10%, 5,780 field plots) datasets.

#### 2.1.3 Landsat imagery

We used Landsat 5 TM, 7 ETM+, and 8 OLI surface reflectance products for predictors of fractional cover. We masked pixels identified as clouds, cloud shadow, snow, and saturated surface reflectance to calculate 64-day means throughout a given year for surface reflectance bands 2-7. The 64-day periods resulted in six measurements per year, with start dates occurring on day of year 001, 065, 129, 193, 257, and 321. To supplement surface reflectance measurements, we calculated normalized difference vegetation index (NDVI) and normalized burn ratio two (NBR2) for each 64-day period. These indices represent vegetation domains that have been successful in modeling rangeland fractional cover (Jones et al., 2018). We reprojected and bilinearly resampled all Landsat imagery to a geographic coordinate system of approximately 30m resolution.

### 2.2 Model

We used Landsat surface reflectance measurements, vegetation indices, and spatial location (XY coordinates) as covariates to predict rangeland fractional cover. To generate a multivariate response, we used a temporal convolutional network to learn and predict cover for each functional group simultaneously. We used a temporal convolution (i.e., one dimensional convolution) as features associated with each vegetation field data plot varied sequentially through time, but not space. Temporal convolutions work well for satellite time series classification (Pelletier, Webb, & Petitjean, 2019; Zhong, Hu, & Zhou, 2019) and may also outperform standard recurrent neural networks such as long short-term memory (Bai, Zico Kolter, & Koltun, 2018).

We combined temporal convolutions with dropout, pooling layers, and fully connected layers (Figure 1). We used an Adam optimizer with a learning rate of 0.0001, a batch size of 32, a convolutional kernel width of three, and a dropout rate of 20% (Srivastava, Hinton, Krizhevsky, Sutskever, & Salakhutdinov, 2014); the number of filters increased from 32 to 128 over three layers, and the dilation rate increased from one to four. We utilized average pooling with a pooling size of 12 to reduce temporal sequences to a single value. We performed convolutions on Landsat surface reflectances and vegetation indices separately due to the differing domains they represent, concatenating layers prior to a fully connected layer (Figure 1). The final layer contained six units, corresponding to the six functional groups. To produce uncertainty estimates, we implemented dropout during prediction (Gal & Ghahramani, 2015), utilizing a 10% dropout rate before the fully connected layer. We repeated predictions four times, averaged results to obtain the predictive output, and calculated variance to estimate uncertainty.

**Figure 1.**
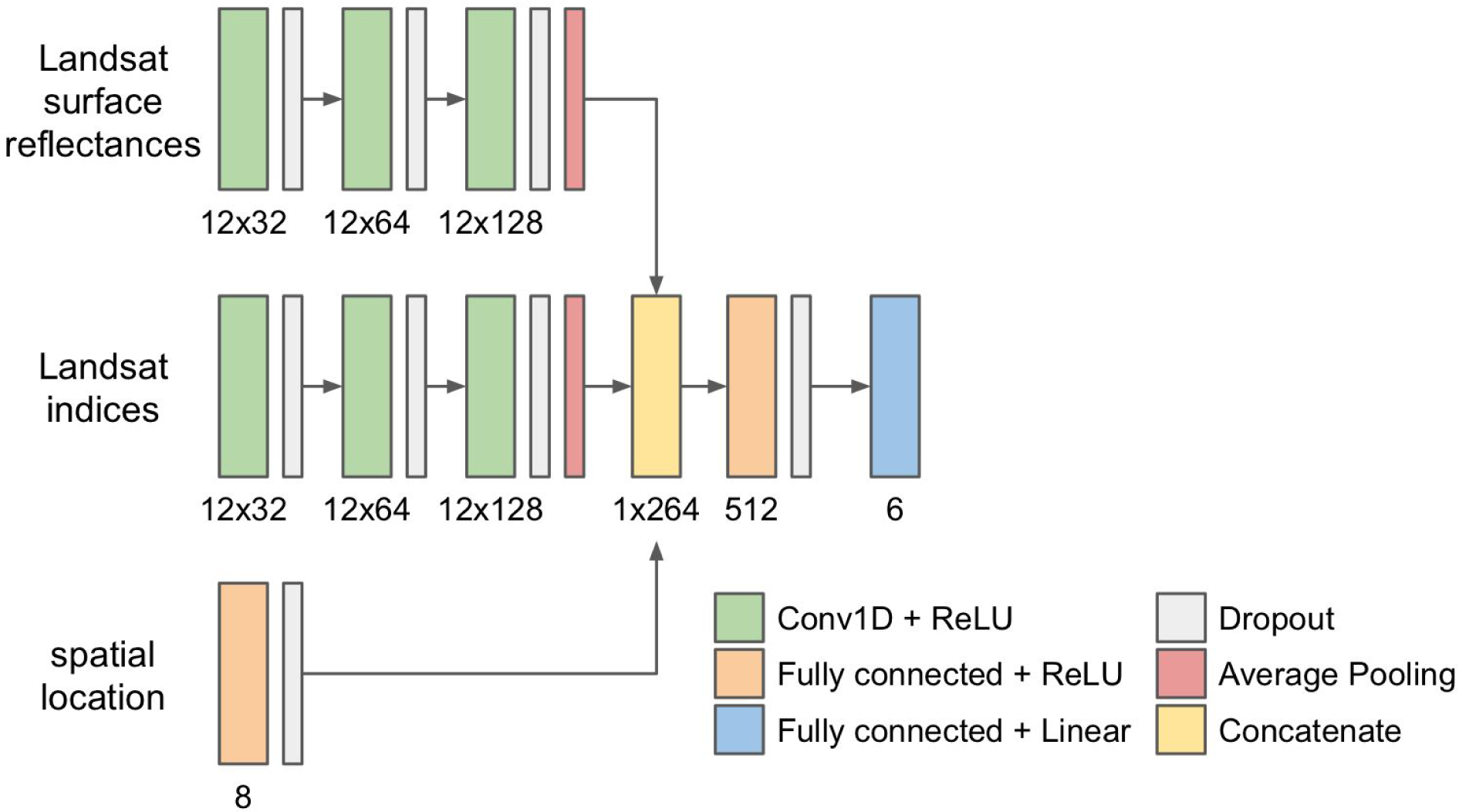
Model architecture to predict cover of rangeland functional groups. Inputs include Landsat surface reflectances, Landsat vegetation indices, and spatial location. Landsat surface reflectances and indices are temporal sequences across twelve 64-day timesteps. The final layer outputs percent cover of the six rangeland functional groups.

We evaluated model performance by calculating mean absolute error (MAE), root mean square error (RMSE), residual standard error, (RSE), and the coefficient of determination (r^2^) of the validation dataset. We compare evaluation metrics to fractional cover version 1.0. In addition to evaluation with the validation dataset, we also evaluated the model with datasets that were not part of the training process and that varied in geographic range, method of collection, scope, and size (Table 1). We developed the model using the Keras library within Tensorflow and performed all image processing and predictions in Google Earth Engine (Gorelick et al., 2017) and Google Cloud AI Platform (AI Platform, 2020), respectively.

**Table 1.** Additional datasets used in evaluation. These datasets varied in geographic range, method of collection, scope, and size.

| Name | Location | Years | n | Description |
| --- | --- | --- | --- | --- |
| Restore New Mexico Collaborative Monitoring Program (RestoreNM) | southwest New Mexico | 2007-2017 | 788 | Plots contained paired parallel 50m transects 20m apart. Transect-level cover used for evaluation. |
| Sagebrush Steppe Treatment Evaluation Project (SageSTEP) | sagebrush steppe | 2006-2014 | 227 | Plots contained 15-24 subplots; five line transects in each subplot. Plot-level cover used for evaluation. |
| Eastern Oregon Agricultural Research Center (EOARC) | eastern Oregon | 2016 | 198 | Plots contained three 20m transects. Plot-level cover used for evaluation. |
| USGS/NPS | Colorado Plateau | 2007-2019 | 3,405 | Plots contained two to three 50m transects. Plot-level cover used for evaluation. See Supporting Information for more information. |
| University of Idaho (UI) | central Idaho | 2018-2019 | 60 | Plots contained three 25m transects. Plot-level cover used for evaluation. |

## 3 Results

### 3.1 Model evaluation

Model results and evaluation metrics suggest strong relationships between predicted and on-the-ground measurements (Table 2 and Figure 2). Evaluation metrics of the validation dataset averaged 6.3 and 9.6% (MAE and RMSE, respectively) across rangeland functional groups. Residual standard errors of predicted and on-the-ground measurements varied from 4.6 to 12.7% among functional groups (Table 2). Coefficient of determination values ranged from 0.57 to 0.77 for most functional groups (Table 2). Evaluation metrics calculated with additional datasets that were not part of the training dataset also revealed similar metrics (Table 3). Model performance increased compared to fractional cover version 1.0 (Jones et al., 2018; Table 2), and is comparable to other US rangeland fractional products available over disparate geographies (Zhang et al., 2019; Rigge et al., 2020).

**Table 2.** Model evaluation metrics (mean absolute error, MAE; root mean square error, RMSE; residual standard error, RSE; and coefficient of determination, r^2^) calculated using the respective validation dataset for fractional cover versions 1.0 and 2.0.

|  | **Annual**<br>**Forb &**<br>**Grass** | **Perennial**<br>**Forb &**<br>**Grass** | **Shrub** | **Tree** | **Litter** | **Bare ground** | **Average** |
| --- | --- | --- | --- | --- | --- | --- | --- |
| **fractional cover version 2.0** (this paper) |  |  |  |  |  |  |  |
| **MAE** | 7.0 | 10.3 | 5.8 | 2.8 | 5.7 | 6.7 | 6.3 |
| **RMSE** | 11.0 | 14.0 | 8.3 | 6.8 | 7.9 | 9.8 | 9.6 |
| **RSE** | 8.8 | 12.7 | 6.6 | 5.9 | 4.6 | 7.9 | - |
| **$r^2$** | 0.58 | 0.77 | 0.57 | 0.65 | 0.37 | 0.73 | - |
| **fractional cover version 1.0** (Jones et al., 2018) |  |  |  |  |  |  |  |
| **MAE** | 7.8 | 11.1 | 6.9 | 4.7 | - | 7.3 | 7.56 |
| **RMSE** | 11.8 | 14.9 | 9.9 | 8.5 | - | 10.6 | 11.14 |
| **RSE** | - | - | - | - | - | - | - |
| **$r^2$** | 0.43 | 0.71 | 0.43 | 0.52 | - | 0.71 | - |

**Table 3.** Model evaluation metrics (mean absolute error, MAE; root mean square error, RMSE; and coefficient of determination, r^2^) calculated using additional datasets described in Table 1.

|  | **Annual**<br>**Forb &**<br>**Grass** | **Perennial**<br>**Forb &**<br>**Grass** | **Shrub** | **Tree** | **Litter** | **Bare ground** |
| --- | --- | --- | --- | --- | --- | --- |
| **RestoreNM** |  |  |  |  |  |  |
| **MAE** | 5.7 | 9.7 | 6.5 | - | - | - |
| **RMSE** | 11.2 | 13.1 | 8.1 | - | - | - |
| **$r^2$** | 0.29 | 0.22 | 0.08 | - | - | - |
| **SageSTEP** |  |  |  |  |  |  |
| **MAE** | 9.0 | 13.2 | 8.3 | - | - | 8.9 |
| **RMSE** | 13.3 | 18.1 | 9.8 | - | - | 11.8 |
| **$r^2$** | 0.19 | 0.25 | 0.27 | - | - | 0.49 |
| **EOARC** |  |  |  |  |  |  |
| **MAE** | 6.6 | 9.5 | 5.8 | - | - | - |
| **RMSE** | 9.6 | 11.8 | 7.6 | - | - | - |
| **$r^2$** | 0.43 | 0.37 | 0.57 | - | - | - |
| **USGS/NPS** |  |  |  |  |  |  |
| **MAE** | 4.2 | 8.5 | 7.6 | 5.3 | 7.41 | 8.1 |
| **RMSE** | 7.8 | 11.7 | 10.32 | 9.6 | 11.2 | 11.1 |
| **$r^2$** | 0.49 | 0.39 | 0.56 | 0.54 | 0.31 | 0.71 |
| **UI** |  |  |  |  |  |  |
| **MAE** | 8.0 | 10.5 | 9.8 | - | - | 3.2 |
| **RMSE** | 10.6 | 13.3 | 12.1 | - | - | 4.3 |
| **$r^2$** | 0.56 | 0.28 | 0.30 | - | - | 0.31 |

**Figure 2.**
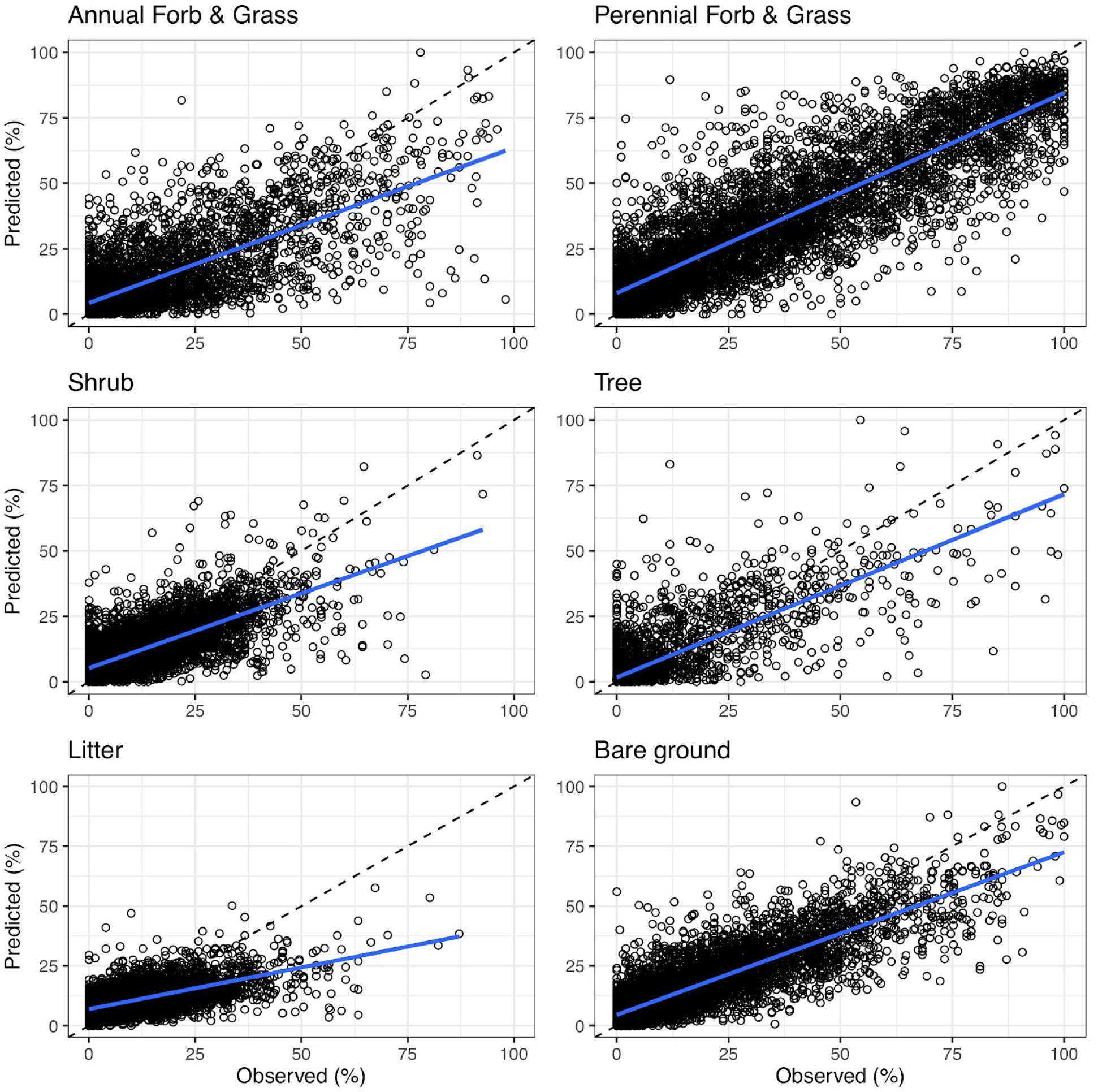
Predictions of fractional cover relative to observed on-the-ground measurements, separated by rangeland functional group. Diagonal dashed black line represents a 1:1 relationship; solid blue line is the linear fit between predicted and observed values. Coefficient of determination (r^2^) and residual standard error (RSE) are reported in Table 2.

## 4 Discussion

We provide next generation predictions of annual, fractional cover of rangeland functional groups by implementing a multitask learning approach across the western US (Figures 3 and 4). We improved upon our previous efforts (Jones et al., 2018) by 1) utilizing a neural network that models the dynamic interactions of functional groups; 2) reducing errors and improving model fit; and 3) providing spatiotemporal-explicit, pixel-level estimates of uncertainty alongside predictions. We deliver these data via the Rangeland Analysis Platform (https://rangelands.app/), an online and interactive web application that tracks rangeland vegetation through time.

**Figure 3.**
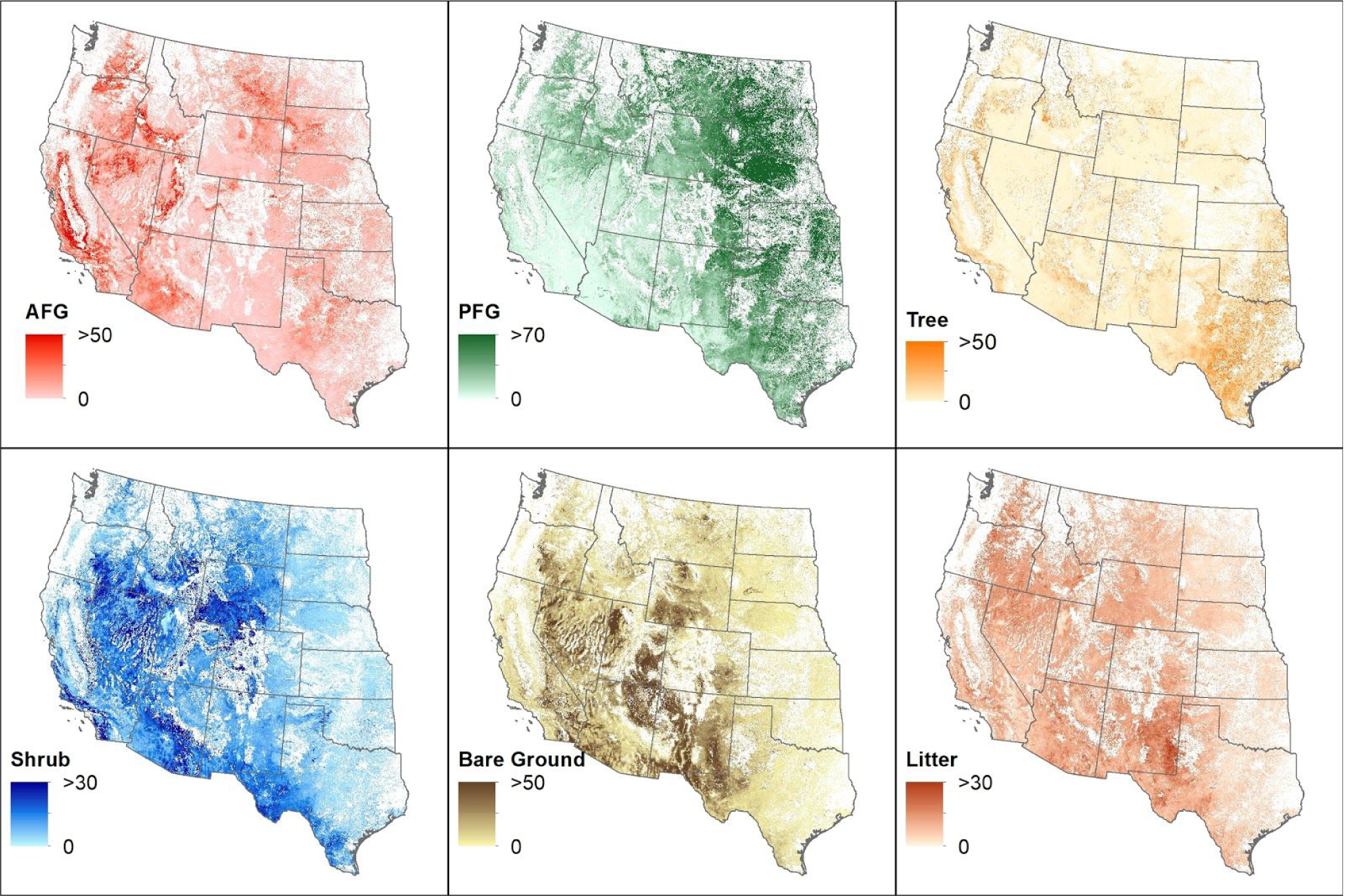
Fractional cover predictions of annual forbs and grasses (AFG), perennial forbs and grasses (PFG), shrubs, trees, litter, and bare ground for 2019. White areas are non-rangeland as identified by Reeves and Mitchell (2011).

**Figure 4.**
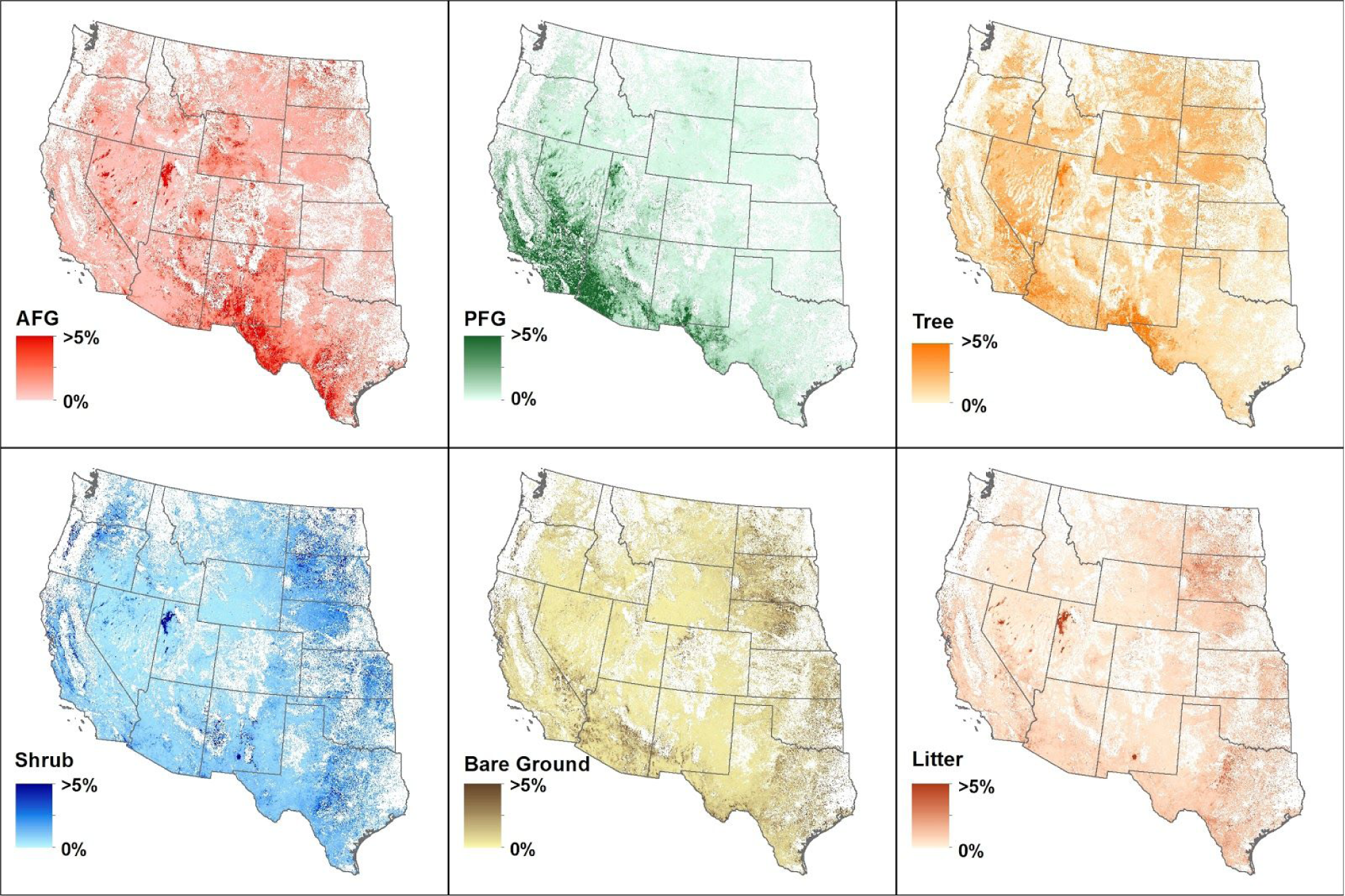
Fractional cover uncertainty of annual forbs and grasses (AFG), perennial forbs and grasses (PFG), shrubs, trees, litter, and bare ground for 2019. White areas are non-rangeland as identified by Reeves and Mitchell (2011). Uncertainty was relativized to the fractional cover prediction.

The fractional cover of functional groups and cover types in rangelands reflect numerous ecosystem processes and ecosystem services. Changes in one functional group has predictable ecological impacts on other groups and the services they provide (Uden et al., 2019). For example, woody plant encroachment into grasslands constrains herbaceous grass cover and diminishes forage production and wildlife habitat (Archer et al., 2017), whereas annual grass invasion reduces perennial herbaceous plants and shrubs (Davies, 2008). While previous univariate modeling methods of fractional cover disregard this inherent covariation, a multitask model exploits it (Caruana, 1997). The underlying relationships among rangeland functional groups are learned and incorporated into the model, increasing accuracy (Figure 5). Although univariate predictions can be constrained or restricted post hoc to correct for or reduce such errors (Henderson, Bell, & Gregory, 2019), the goal of multitask learning is to learn and predict variables simultaneously. Furthermore, the shared representation of multitask learning allows for covariance dynamics and interactions to be defined by the data, eliminating the need for predetermined conditions, rules, or thresholds.

**Figure 5.**
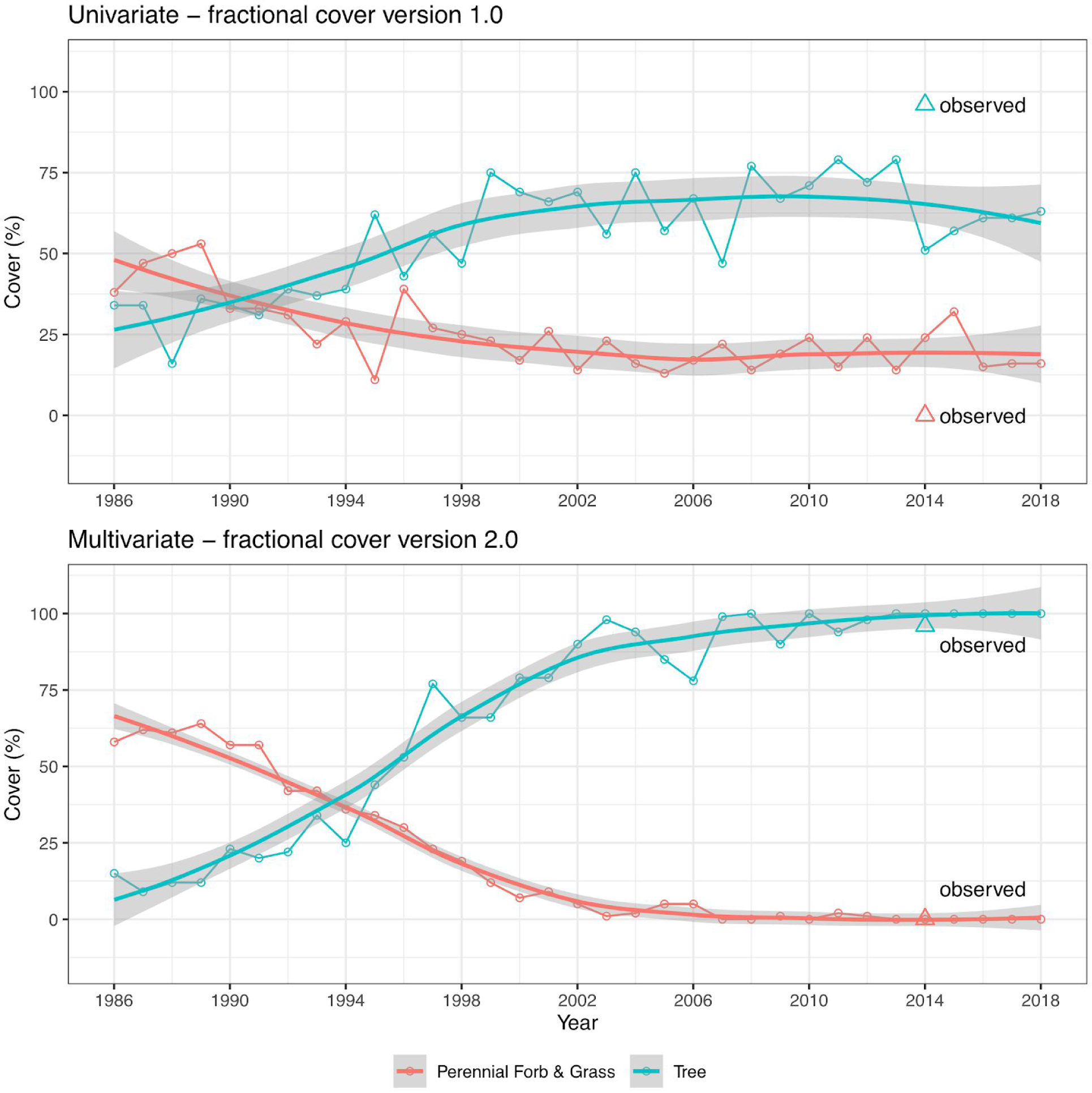
Univariate (top) and multivariate (bottom) predictions of perennial forb/grass and tree fractional cover for a single validation plot in an area with woody encroachment. Plot data was collected in 2014 and recorded 0 and 96% cover (triangular points) for perennial forb/grass and tree, respectively. Due to shared representation, multitask models and predictions better represent functional group dynamics. Fractional cover version 1.0 produced by Jones et al. (2018). Shaded lines represent locally estimated scatterplot smoothing.

When developing remotely sensed products, the goal is often to maximize model performance to increase accuracy. Error, however, is always present and should be understood and integrated into the application of that product and the decision being informed. Common model error metrics measure the average difference between predicted model output and individual on-the-ground measurements. This is generally calculated by withholding a portion (e.g., 5-20%) of the model training dataset for validation (either entirely, or in a bootstrap aggregating approach). Error metrics therefore represent an average accuracy for the model given the validation dataset, but do not indicate any spatial or temporal variability of error. Attempts to visualize or aggregate errors across broad regions may appear helpful, but do little to characterize their spatial distribution or to help judge spatial accuracy (Jones et al., 2018; Zhang et al., 2019). Moreover, error is commonly calculated with only a minuscule fraction (<<0.1%) of known measurements relative to total predictions, e.g., the 5,780 known measurements used for validation (10% of the training dataset) in this exercise represents approximately 2e-8% of total predictions.

The advantage of uncertainty information over error is its spatiotemporal utility: uncertainty is calculated with every prediction and therefore varies across space and time (Figure 6), while error does not (see discussion above). Uncertainty provides a measure of prediction confidence, i.e., how reasonable is this prediction given the data used to build the model? If the characteristics of a location are generally represented within the model training data, uncertainty may be low and the corresponding prediction reasonable. If not as well represented within the model training data (e.g, areas such as high alpine, lava flows, arid playas), uncertainty may be high and the corresponding prediction unreasonable. Due to the fact that uncertainty is spatiotemporally explicit, it can be helpful in determining how to use model predictions on a case-by-case basis. For example, if uncertainty is high in a particular area or time period of interest, a practitioner can choose to gather additional local data or information, do a more detailed analysis, discuss with colleagues, etc. in order to expand decision inputs. Uncertainty can be integrated into the context of the decision being made for a particular place or time.

**Figure 6.**
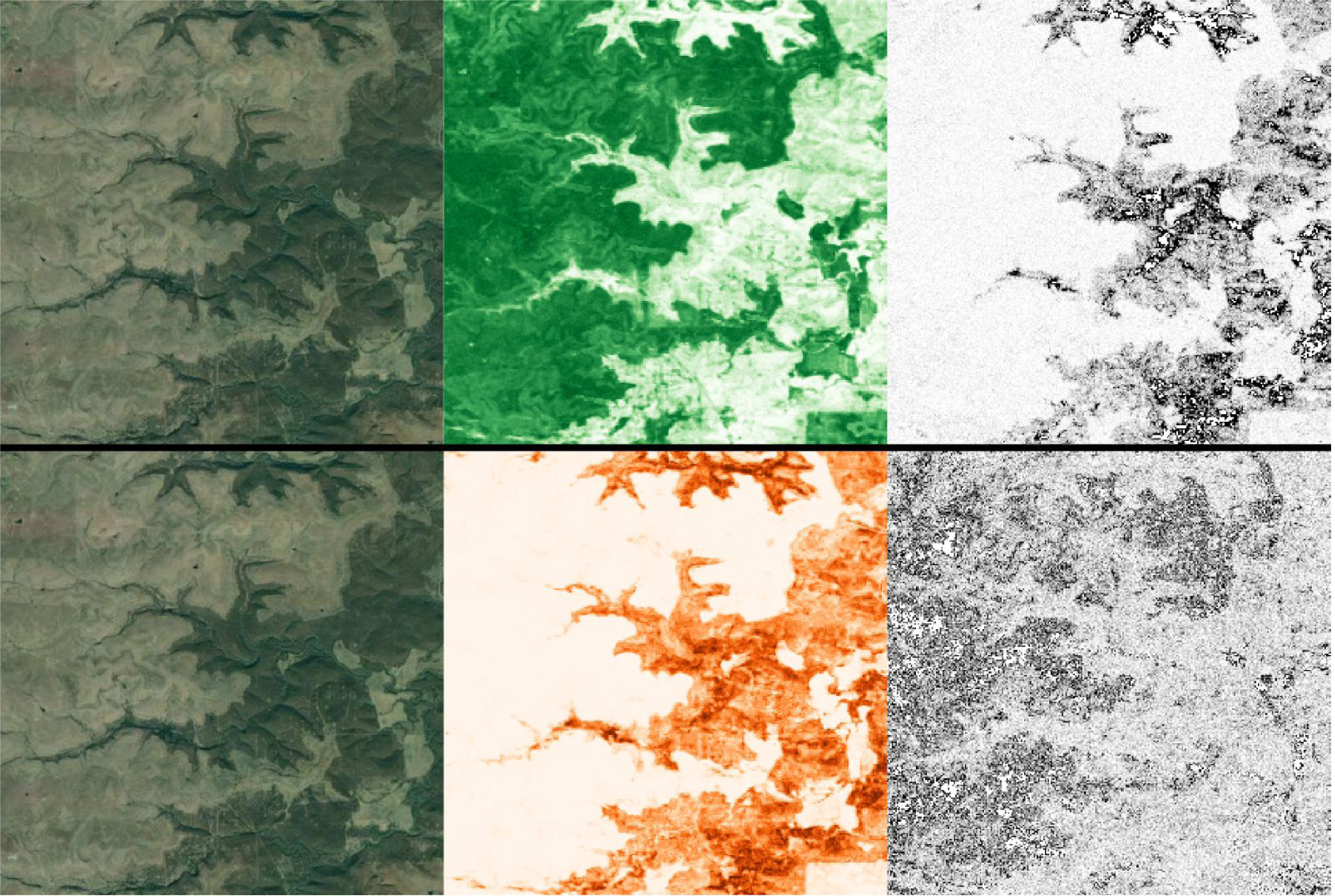
Aerial imagery (left), cover estimates (middle), and uncertainty estimates (right) for 2019 perennial forb and grass (top) and tree (bottom) functional group cover for a small region in the southern Great Plains. Light-to-dark values represent lower-to-higher values of cover and uncertainty. Greater uncertainty for perennial forb and grass estimates are present in areas dominated by trees and vice versa.

Operationalizing uncertainty information into decision making presents new opportunities to learn about this type of information. While users often simply want to know if a prediction is “right or wrong” or “how far off it is”, it is important to note that uncertainty does not provide this. Rather, uncertainty is a measure of model confidence (more specifically prediction variance), and should be thought about, processed, and utilized in the same way that other confidence-, odds-, or probability-type information is consumed, e.g., precipitation probabilities supplied with weather forecasts. As such, there are no defined rules, standardized practices, thresholds, etc. to immediately employ when using uncertainty information. Use of such methods will vary depending upon the context of the decision being made. We recognize that it may be difficult to immediately incorporate uncertainty information into decision making frameworks and workflows. We are confident, however, that with increased education, experience, and exposure to these types of information, such barriers will be lessened and removed.

## 5 Conclusions

Innovations in remotely sensed mapping of rangeland cover continue to present new opportunities to improve assessment, management, and monitoring. We provide the latest advancement to expand the land management toolbox with improved predictions of fractional cover at a moderate resolution of 30m, along with spatiotemporal-explicit uncertainty estimates, that can be used at such resolution or aggregated to broader scales. This product is intended to be used in combination with local on-the-ground data, expert knowledge, land use history, scientific literature, and other sources of information when making interpretations. We emphasize that when being used to inform decision-making, remotely sensed products should be evaluated and utilized according to the context of the decision and not be used in isolation. Learning how to think about and use remotely sensed data, and suitably integrate them into decision frameworks and workflows, are next steps for improving the field of rangeland monitoring.

## Acknowledgements

We thank Anna Knight and Michael Duniway for assistance with data evaluation.

## Author contributions

BA, MJ, JM, DN, TG, and DT led the writing of the manuscript. BA, CB, TE, MJ, JM, SM, and DU conceived, designed, implemented, or assisted with methodology. BB, CB, KD, MD, LE, SF, VJ, JK, AK, and HS collected, provided, or analysed data for additional evaluation. All authors contributed to drafts and gave final approval for publication.

## Data availability

Data are available on the Rangeland Analysis Platform (https://rangelands.app/) and from http://rangeland.ntsg.umt.edu/data/rap/rap-vegetation-cover/.

