## Supporting information for "Improving Landsat predictions of rangeland fractional cover with multitask learning and uncertainty"

### USGS/NPS data description

Field-sampled validation plots from across the Colorado Plateau have been compiled and curated by the US Geological Survey Southwest Biological Science Center in Moab, UT (Fig. S1). Data sources include monitoring data from the National Park Service Inventory and Monitoring - Northern Colorado Plateau Network (I&M-NCPN, n = 2429; Witwicki et al., 2017), plots from the Bureau of Land Management Canyon Country District Office fuels program (BLM-CCDO, n=404), and several projects run by the US Geological Survey in Moab, UT (USGS-Moab, n=572; Duniway and Palmquist, 2020; Duniway et al., 2015; Fick et al., 2020; Miller et al., 2012; Van Scoyoc, 2014; Webb et al., 2016). In total, 3,405 plot sampling events were used. All data for the indicators used for this validation effort were collected using the line-point intercept method (Herrick et al., 2009). Plots consisted of two to three 50-m transects. All I&M-NCPN plots and most USGS-Moab plots consisted of three parallel transects, spaced 25 m apart. Some plots in the USGS-Moab and BLM-CCDO data sets had transects arranged in a hub-and-spoke pattern. Indicators were calculated using the terradactyl package in R (in development by Sarah McCord and Stauffer, 2020). All indicators use first hit to calculate cover. Plots were sampled between May 2007 and June 2019.

Rangeland Analysis Platform (RAP) 2.0 predictions were extracted within a 35-m radius of the plot center point to completely encompass all transects in plots with parallel transects. For plots sampled more than once, each field sampling event was compared with RAP predictions independently. Fig. S2 compares field measurement to RAP predictions of six indicators.

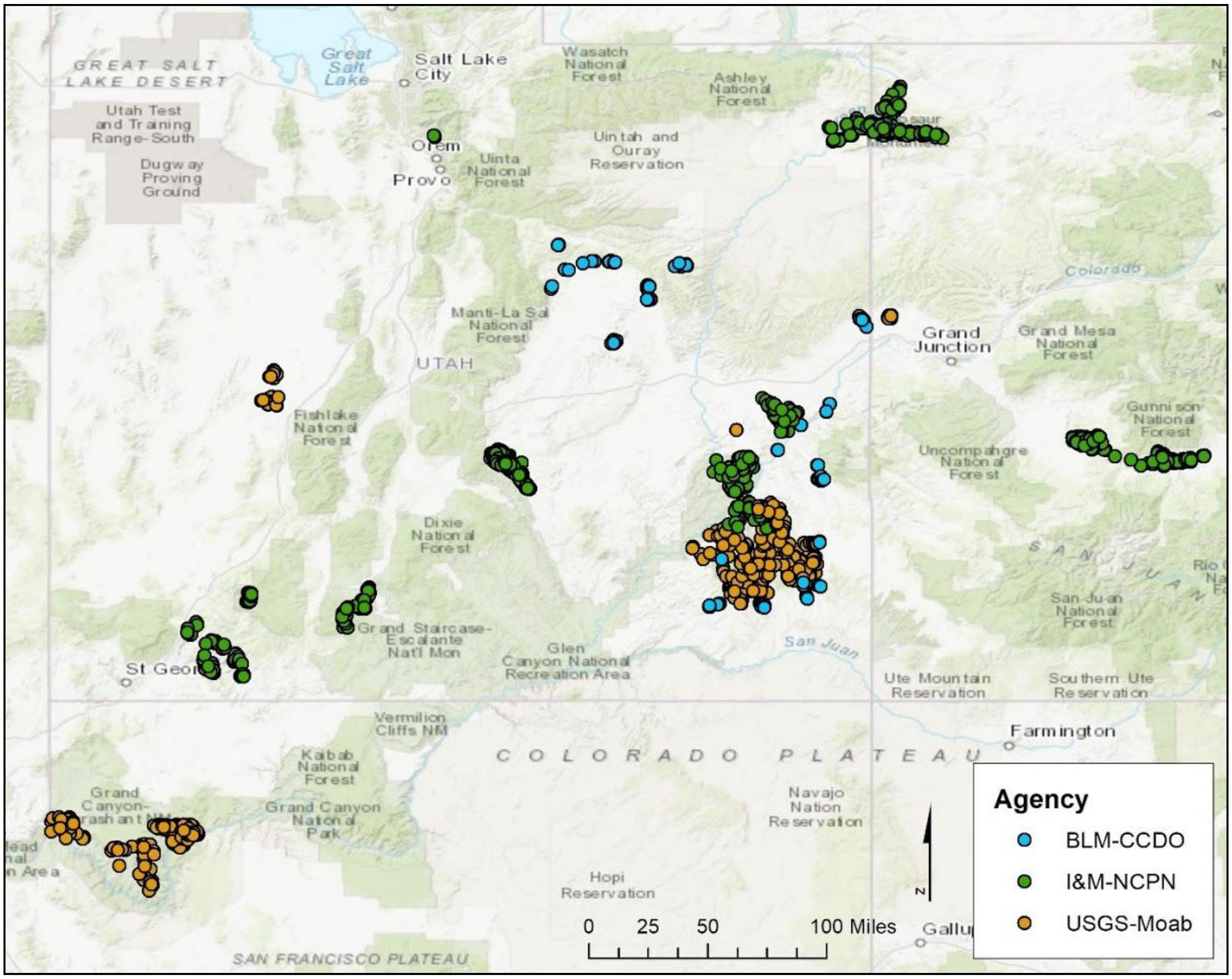

Figure S1. Field-sampled plot locations on the Colorado Plateau.

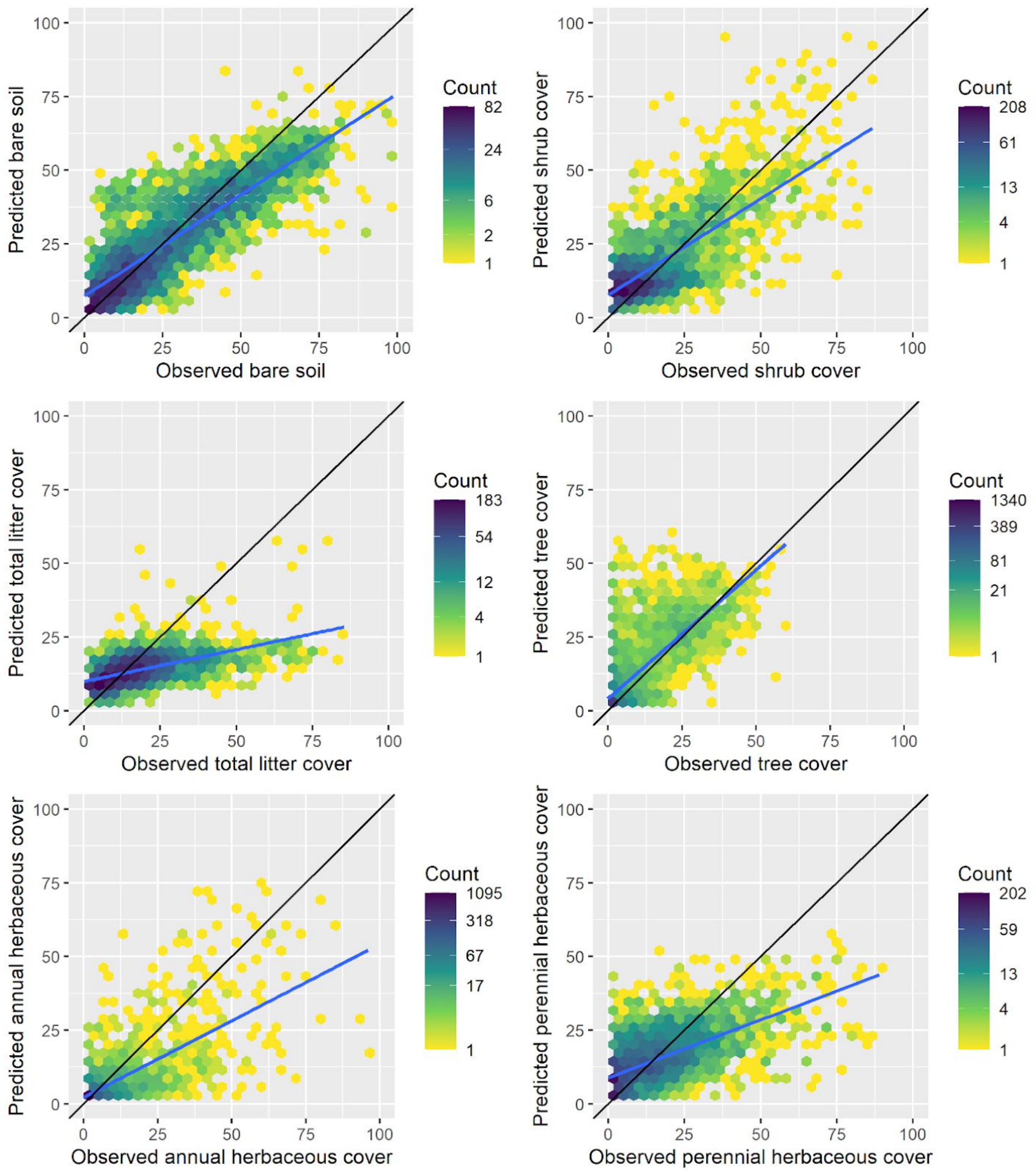


Figure S2. Field sampled indicators (observed) vs. RAP indicators (predicted). The blue line is a linear model of field vs. RAP 2.0 values; the black line is a 1-1 line.
